# RiboFlow, RiboR and RiboPy: An ecosystem for analyzing ribosome profiling data at read length resolution

**DOI:** 10.1101/855445

**Authors:** Hakan Ozadam, Michael Geng, Can Cenik

## Abstract

**Summary:** Ribosome occupancy measurements enable protein abundance estimation and infer mechanisms of translation. Recent studies have revealed that sequence read lengths in ribosome profiling data are highly variable and carry critical information. Consequently, data analyses require the computation and storage of multiple metrics for a wide range of ribosome footprint lengths. We developed a software ecosystem including a new efficient binary file format named ‘ribo’. Ribo files store all essential data grouped by ribosome footprint lengths. Users can assemble ribo files using our RiboFlow pipeline that processes raw ribosomal profiling sequencing data. RiboFlow is highly portable and customizable across a large number of computational environments with built-in capabilities for parallelization. We also developed interfaces for writing and reading ribo files in the R (RiboR) and Python (RiboPy) environments. Using RiboR and RiboPy, users can efficiently access ribosome profiling quality control metrics, generate essential plots, and carry out analyses. Altogether, these components create a complete software ecosystem for researchers to study translation through ribosome profiling.

**Availability and Implementation:** For a quickstart, please see https://ribosomeprofiling.github.io. Source code, installation instructions and links to documentation are available on GitHub: https://github.com/ribosomeprofiling

## Introduction

Ribosome profiling is a powerful method for measuring transcriptome-wide translation through the sequencing of ribosome-protected mRNA fragments (Ingolia *et al.*, 2009, 2011). This transformative approach has been applied to many organisms and can approximate translational efficiency, a major determinant of protein abundance (Schwanhäusser *et al.*, 2011; Kristensen *et al.*, 2013). Consequently, ribosome profiling studies fulfill a critical gap in our understanding of protein abundance, which is only partially explained by RNA expression (Marguerat *et al.*, 2012; Ly *et al.*, 2014). Moreover, ribosome profiling has proved invaluable in studying the mechanisms of translation (Ingolia *et al.*, 2019).

Initial ribosome profiling studies focused on the classical ∼28 nt ribosome-protected footprints (RPFs) corresponding to ribosomes with occupied A-sites. In contrast, recent work revealed that RPF lengths are variable and carry critical information (Lareau *et al.*, 2014; Wu *et al.*, 2019; Liakath-Ali *et al.*, 2018). For example, ribosomes can protect short footprints (15-21 nts), characteristic of different ribosome conformations (Guydosh and Green, 2017; Lareau *et al.*, 2014; Wu *et al.*, 2019), as well as longer footprints (∼60 nt) indicative of ribosome collisions (Arpat *et al.*; Guydosh and Green, 2014). Importantly, these key observations signal a new chapter of translation studies that will need to be analyzed by taking variable RPF lengths into account.

Analyses for a wide range of RPF lengths (∼15-60) entail increased computational complexity and organizational challenges that were negligible in ribosome profiling studies that focused solely on ∼28 nt RPFs. In particular, the naive use of text files would be highly inefficient in organization, computation, and storage. Similarly, directly accessing the unprocessed alignment files for analyses limits data portability and efficiency. While a range of computational approaches have been developed for ribosome profiling, they either focus solely on ∼28 nt footprints (Berg *et al.*, 2019; Popa *et al.*, 2016), or directly rely on alignment files for analyses and do not consider RPF length as a major design feature (Carja *et al.*, 2017; Chung *et al.*, 2015; Perkins *et al.*, 2019; Birkeland *et al.*, 2018). Moreover, existing software often lags behind high-quality standards in installation, ease of use, documentation, and portability (Wang *et al.*, 2019).

We introduce a new binary file format called ‘ribo’ to enable efficient organization of ribosome profiling data including studies focusing on a broad range of RPF lengths. Similar binary formats are widely-used to store many sequencing types such as ‘bam’ for sequence alignments (Li *et al.*, 2009), ‘BUS’ for single-cell RNA-Seq (Melsted *et al.*, 2019), and ‘cooler’ / ‘hic’ for Hi-C (Abdennur and Mirny, 2019; Durand *et al.*, 2016). Our new ribo file is designed to work with ribosome profiling data at nucleotide length resolution. Ribo files organize all quantification tables and metadata for efficient data storage and retrieval.

Importantly, we designed a software ecosystem around this file format that includes three major components. RiboFlow is a Nextflow (Di Tommaso *et al.*, 2017) based alignment pipeline that generates ribo files from raw sequencing data. Riboflow can run on local servers, major job schedulers, and cloud based platforms with minimal configuration. We provide a Docker container image to enable deployment of RiboFlow on all major operating systems. Finally, we developed two interfaces, RiboPy and RiboR, to work with ribo files. RiboPy, a Python package, can be used to create ribo files and subsequently analyze and visualize data. RiboR is a package that enables seamless analyses with ribo files in the R environment. Importantly, we offer a superior user experience facilitating data portability, enabling installation via package management tools and providing detailed documentation. Taken as a whole, this ecosystem is a complete platform for researchers to study translation using ribosome profiling.

## Implementation and Availability

### RiboFlow

RiboFlow is a Nextflow (Di Tommaso *et al.*, 2017) based pipeline that generates ribo files from sequencing files in one step. The RiboFlow pipeline begins with adapter removal from raw sequencing reads in fastq format. The clipped reads are then filtered to remove common noncoding RNAs such as ribosomal RNA. Next, remaining reads are mapped to the transcriptome, and alignments with a mapping quality higher than a predetermined value are retained. An optional PCR deduplication step can collapse PCR duplicates defined by having the same read length and identical mapping position. Finally, results from multiple experiments are compiled into one ribo file (Figure 1A).

**Figure 1:**
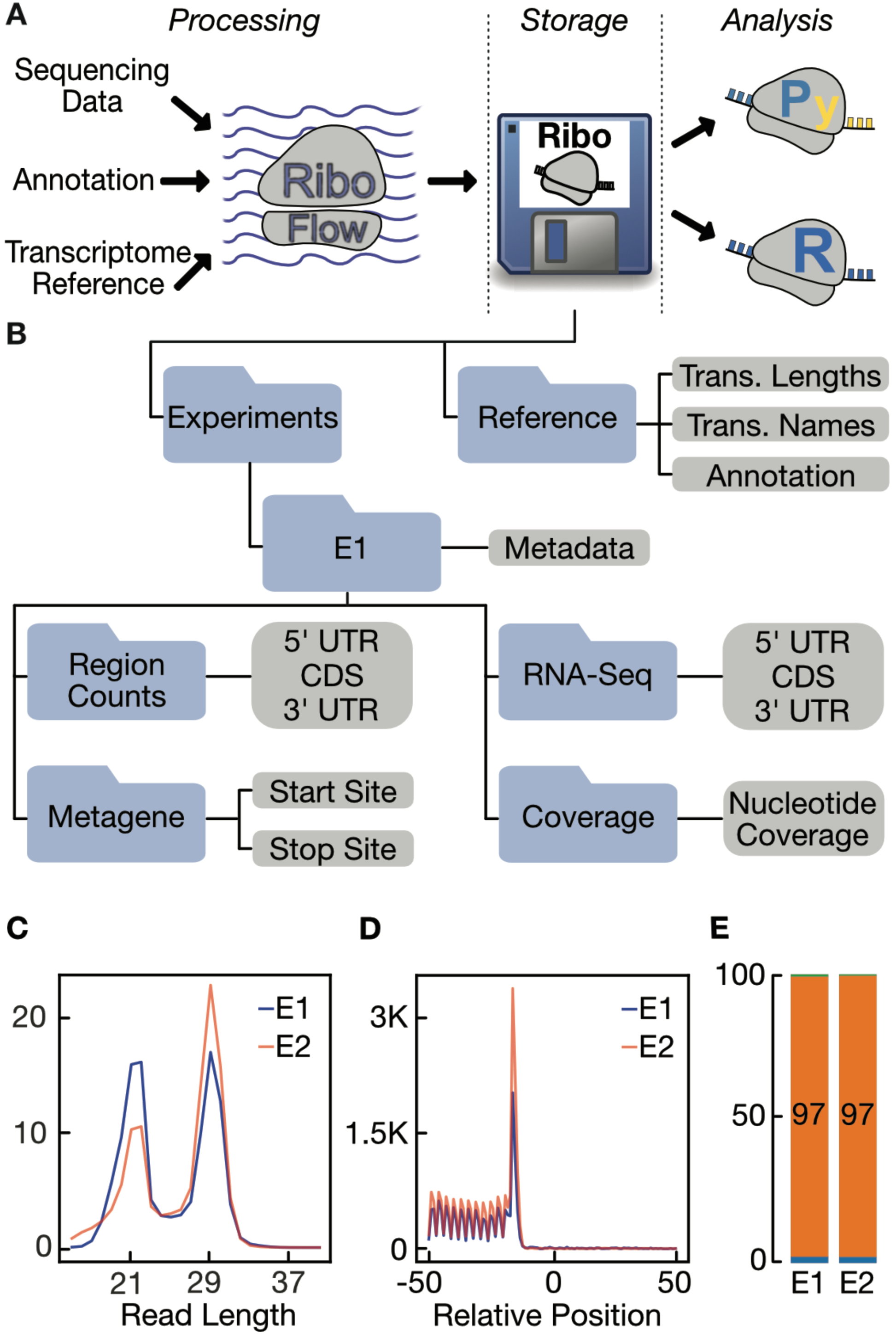
(A) Overview of the Ribo Software Ecosystem. (B) The internal structure of a ribo file is depicted. (C, D, E) are example plots generated by RiboPy from two experiments denoted as E1 and E2. (C) Read length distribution of coding region mapping reads were plotted. The y-axis corresponds to the percentage of each read length. (D) RPFs mapping to the vicinity of the translation stop site are aggregated across genes to display a metagene plot. The y-axis corresponds to reads per million. (E) Percentage of reads mapping to different transcript regions are shown. Top (green) and bottom (blue) regions account for a small fraction of the data as they represent the 3’ and 5’ UTR mapping RPFs, respectively. Middle bar (orange) depicts the percentage of CDS mapping RPFs.

Using the provided Docker container image, RiboFlow can run on all major operating systems. The number of simultaneous threads can easily be set, allowing for efficient hardware utilization across a wide spectrum of computational resources. RiboFlow can be configured to run on mainstream job schedulers such as LSF, SGE, and SLURM as well as cloud environments such as Amazon Web Services and Google Cloud Platform.

### Ribo File

The output of RiboFlow is a binary file named ‘ribo’ which is built on top of the Hierarchical Data Format (HDF) (The HDF Group, 1997-2019). Every ribo file must contain three types of data (Figure 1B):

1. **Transcriptome Annotation:** For each transcript in a given transcriptome, a ribo file contains their names, lengths, and region annotations (5’ UTR, CDS, 3’ UTR).
2. **Region RPF Counts:** Ribosome profiling data is quantified using the number of reads mapping to the different transcript regions, namely the 5’UTR, CDS, and 3’UTR. The distribution of reads across these regions can be informative as a quality control metric and depend on the RNase of choice in the experiment (Miettinen and Björklund, 2015; Wolin and Walter, 1988).
3. **Metagene Counts:** Read counts are aggregated across all transcripts with respect to the translation start/stop sites. This data summarization is typically referred to as ‘metagene counts’.

The following data are optionally included in a ribo file:

#### RNA-Seq Quantification

Most ribosome profiling experiments employ matched total RNA sequencing (RNA-Seq) to enable analyses of translation efficiency. RNA-seq quantifications are stored in a manner that parallels the region counts for the ribosome profiling experiment.

#### Coverage Data

Coverage data refers to the number of reads whose 5’ ends map to each nucleotide position for every transcript.

#### Metadata

A ribo file may contain metadata for any experiment or for the entire ribo file itself. Metadata is defined by the user on a key-value basis.

As a case study, we processed ∼7.3 billion reads across 58 ribosome profiling experiments from three studies (Cenik *et al.*, 2015; Sidrauski *et al.*, 2015; Wu *et al.*, 2019) (GEO accession numbers: GSE65912, GSE65778, GSE115162, respectively). For all three datasets, ribo files yielded more than 15-fold reduction in file size compared to gzip compressed text files (see supplementary material). For example, a ribo file containing all the above-described data from 50 experiments generated in (Cenik *et al.*, 2015) was only 110 MB in contrast to 1711 MB when using gzip compressed text files.

### RiboR and RiboPy

To interact with ribo files, we offer an R package (RiboR) and a Python interface (RiboPy). RiboR and RiboPy provide a set of functions for data import into an R or Python environment and for commonly used visualization. In one function call, users can read ribosome occupancy around the start or stop sites (metagene data), or total read counts for a given transcript region (5’ UTR, CDS, or 3’ UTR). Data can be aggregated or accessed for each individual transcript given a range of RPF lengths. Optional data such as metadata, transcript abundance and ribosome occupancy at nucleotide resolution can be obtained in a similar fashion. Finally, the built-in functions can generate visualizations for RPF length distribution, metagene data, and region-specific read counts (Figure 1, panels C,D and E).

## Conclusions

We describe the first specialized binary file format designed for ribosome profiling data. Storing data in ribo files not only reduces file sizes significantly but also facilitates efficient organization, portability, and data analyses. We developed two interfaces, RiboR and RiboPy, for the most commonly used programming languages in bioinformatics, R and Python. RiboFlow allows users to process their raw sequencing data to generate ribo files across a wide spectrum of computational platforms, ranging from personal computers to high performance computing clusters. For further convenience and reproducibility, software installation and management is handled by package managers and container images. Given those features, this ecosystem offers a complete solution to study ribosome profiling data and provides superior user experience.

## Supplementary Material

All supplementary material is available at https://github.com/ribosomeprofiling/ribo_manuscript_supplemental.

## Acknowledgements

This work was supported in part by NIH grant CA204522 (CC). Can Cenik is a CPRIT Scholar in Cancer Research supported by CPRIT Grant RR180042. The authors acknowledge the Texas Advanced Computing Center (TACC) at The University of Texas at Austin for providing high performance computing and storage resources that have contributed to the research results reported within this paper. URL: http://www.tacc.utexas.edu. We also would like to thank Muxin Wang for her support in building filter sequence files, Xiaoying Wei for software testing and generation of annotation files for mouse, and William Zhang for software testing and processing raw data from Cenik et al. 2015.

## Author Contributions

Can Cenik - Conceptualization, Funding acquisition, Methodology, Project administration, Software Testing, Supervision, Writing - original draft, Writing - review & editing

Michael Geng - Software for RiboR (design, documentation, implementation, testing, vignette drafting), Writing - review & editing

Hakan Ozadam - Methodology, Project administration, Software, Supervision, Validation, Visualization, Writing - original draft, Writing - review & editing

## Notes

https://github.com/ribosomeprofiling

https://ribosomeprofiling.github.io

https://github.com/ribosomeprofiling/ribo_manuscript_supplemental

